# Neural integration and segregation revealed by a joint time-vertex connectome spectral analysis

**DOI:** 10.1101/2022.07.26.501543

**Authors:** Joan Rué-Queralt, Valentina Mancini, Vincent Rochas, Caren Latrèche, Peter J Uhlhaas, Christoph M. Michel, Gijs Plomp, Stephan Eliez, Patric Hagmann

## Abstract

Brain oscillations are produced by the coordinated activity of large groups of neurons and different rhythms are thought to reflect different modes of information processing. These modes, in turn, are known to occur at different spatial scales. Nevertheless, how these rhythms support different modes of information processing at the brain scale is not yet fully understood. Here we present “Joint Time-Vertex Connectome Spectral Analysis”, a framework for characterizing the spectral content of brain activity both in time (temporal frequencies) and in space (spatial connectome harmonics). This method allows us to estimate the contribution of integration (global communication) and segregation (functional specialization) mechanisms at different temporal frequency bands in source-reconstructed M/EEG signals, thus providing a better understanding of the complex interplay between different information processing modes. We validated our method on two different datasets, an auditory steady-state response (ASSR) and a visual grating task. Our results suggest that different information processing mechanisms are carried out at different frequency channels: while integration seems to be a specific mechanism occurring at low temporal frequencies (alpha and theta), segregation is only observed at higher temporal frequencies (high and low gamma). Crucially, the estimated contribution of the integration and segregation mechanisms predicts performance in a behavioral task, demonstrating the neurophysiological relevance of this new framework.

## Introduction

Brain oscillations are produced by the coordinated activity of large groups of neurons and are interpreted as intermittent changes in neuronal excitability (1). Human brain oscillations can be measured at the scalp by electroencephalography (EEG). Traditional approaches to EEG analysis involve the decomposition of the EEG signal into functionally distinct frequency bands, each of which has different temporal and topographical characteristics. In adults, typical frequency bands and their approximate spectral boundaries are delta (1–3 Hz), theta (4–7 Hz), alpha (8–12 Hz), beta (13–30 Hz), and gamma (30–100 Hz) (1,2).

Distinct brain rhythms are thought to support different modes of information processing. Theta oscillations have been proposed to subserve the attentional sampling of the incoming sensory stimuli (3). Alpha and beta oscillations are thought to mediate top-down attentional mechanisms (3), while gamma-band oscillations are thought to reflect the propagation of incoming stimuli in the respective primary sensory cortex (3). In addition, neurophysiological studies revealed the specificity of cortical layers in the visual system. Theta and gamma rhythms originates from the supragranular layers and serve as feedforward signaling, while feedback influences are carried by alpha and beta band oscillations which arise from infragranular layers (4,5).

For these reasons, brain oscillations have emerged as an important feature of large-scale networks, with the potential to link specific patterns of neural activity to behavior (1). Recently, it has been shown that the oscillatory activity in distinct frequency bands is differentially constrained by the anatomical architecture of the white matter, i.e., the connectome in primates (6). However, how distinct brain rhythms support different modes of information processing at the whole-brain level in humans remains unknown.

At the whole-brain level, integration and segregation have been described as the two basic mechanisms of information processing (7). The healthy human brain segregates and integrates information from sensory inputs, proprioception and memories (8). Integration is a global communication mechanism, combining information from different sources to predict the future and adapt behavior accordingly. Segregation is a functional specialization mechanism, in which information is relayed to localized specialized brain regions. One possible way to study the mechanisms underlying segregation and integration is to use neuroimaging methods to map the structure and function of the whole-brain (9), i.e., the spatial and temporal aspects of brain computations.

According to communication through coherence (CCT) account (3,10), the transmission of information between distant neuronal populations is mediated by different oscillatory frequencies. High-frequency oscillations modulate short-range local influences, while long-distance interactions are driven by slower rhythms (11). This property falls within the scope of the standing wave model, which explains the generation of wave patterns in dynamical systems (12). According to this model, wave patterns arise from a direct association between the spatial structure and its temporal frequency of oscillation. A priori, all physical objects have a set of preferred frequencies (*natural frequencies)* at which the system tends to oscillate. Intuitively, one can refer to the oscillations of a violin string, where the typical (resonant) frequencies of oscillations are determined by the length of the string; the longer the string, the lower the fundamental frequency. Similarly, intracortical ECoG studies have shown that, for distant electrodes in the brain, the correlation between the oscillatory activity is higher for low-frequency oscillations than for high-frequency oscillations (13–15), and vice-versa. This suggests that, whereas low-frequency oscillations are better suited to modulate the amplitude of neural activity over distant brain regions in longer temporal windows, high-frequency oscillations modulate activity over neighboring brain regions in shorter temporal windows (16).

Novel tools derived from the field of graph signal processing (12) allow us to estimate the natural normal modes (spatial frequencies) of the connectome, the so-called *connectome harmonics* (Fig. 1c). Connectome harmonics are expected to be the preferred frequencies at which the system tends to enter resonance (17), and have been proposed to capture the two main modes of neural information processing, namely segregation and integration (18–20). Low-frequency connectome harmonics correspond to patterns of activity associated with long-range functional connectivity, with a smooth change in value over distant nodes (17). In contrast, high-frequency connectome harmonics correspond to short-range connectivity patterns, i.e., only brain regions that are directly (and strongly) connected show similar values (Fig. 1d). Consequently, it has been suggested that low-frequency harmonics reflect integration mechanisms, whereas high-frequency harmonics capture segregation mechanisms (18–20) (Fig. 1d).

**Figure 1.**
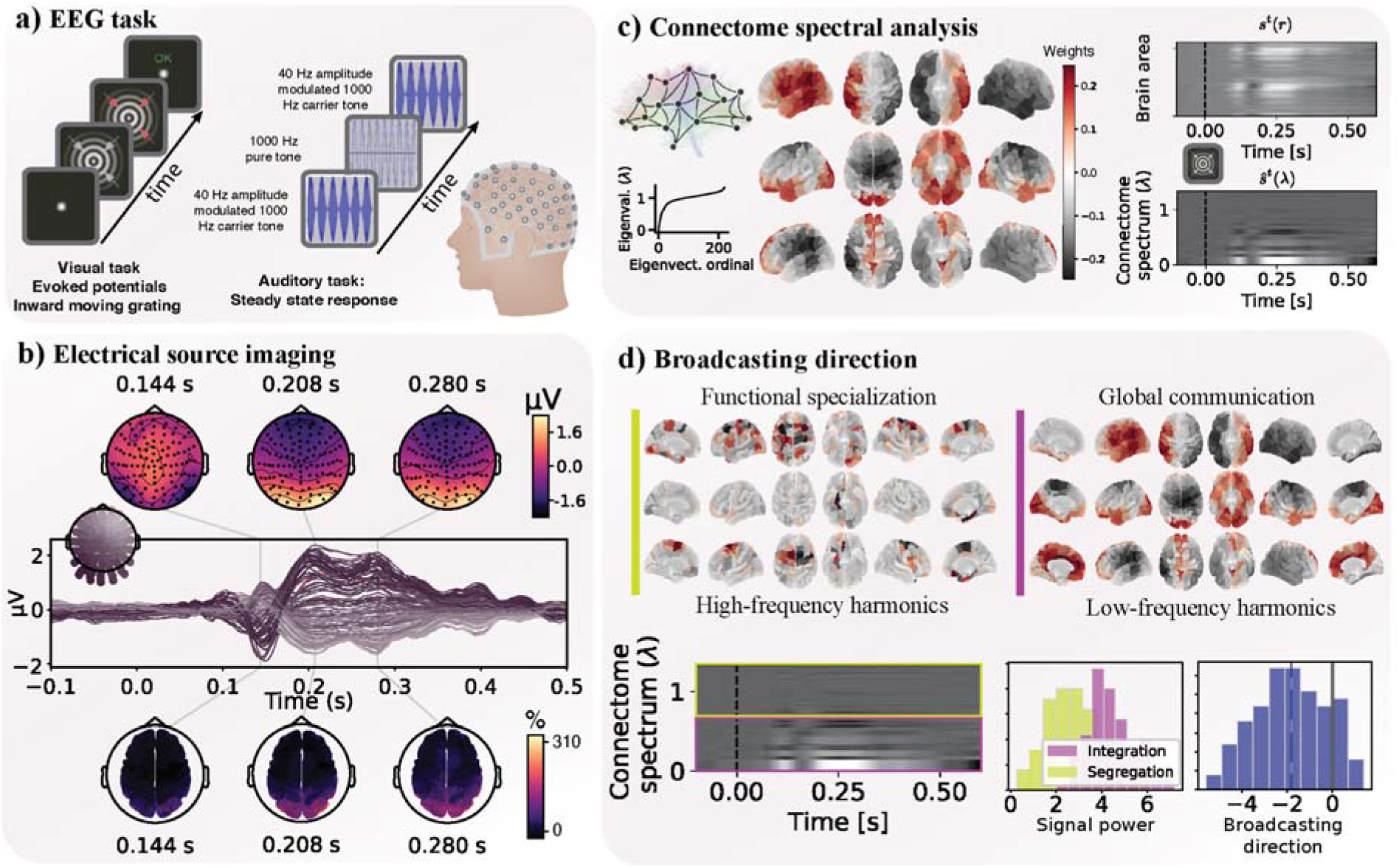
Methodological pipeline. **a) Description of the EEG tasks**. Left: diagram of the visual inward moving grating task. Participants are asked to report the change in speed of inward motion of the grating by button press. Right: 40 Hz Auditory steady state response (ASSR) paradigm composed by 100 ripple tones and 10 intermixed flat tones, each lasting 2 seconds. Participants were asked to recognize flat tones only responding by button press. **b) Electrical source imaging**. Scalp patterns before, and after the sensory stimuli are presented (at 0 ms) and their source reconstruction at three time-points. **c**) **Connectome spectral analysis**: gray matter areas defined by an anatomical parcellation define the nodes of the brain graph, and the estimated white matter tracts from <u>diffusion MRI</u> define the connectivity strength of the edges of the graph. From the constructed graph, we perform the graph Laplacian eigendecomposition and obtain a graph spectrum, consisting of a set of eigenvalues and a set of eigenvectors, the former ones termed graph spectrum, and the later termed graph or connectome harmonics. The source reconstructed time-series signal represented in each brain area (top plot) or in each connectome harmonic (bottom plot). **d**) **Broadcasting direction**: High-(low-)frequency connectome harmonics capture the patterns of segregation (integration) mechanisms. The difference between the power in each of these types of mechanisms defines the broadcasting direction.

When first discovered, connectome harmonics were shown to form the building blocks of well-known functional brain networks associated with both resting-state activity and different cognitive tasks (17). Recent work suggests that connectome harmonics can predict the specific temporal frequency range of a connectivity network in both functional MRI (21) and MEG (22) recordings, indicating that the temporal synchronization between neural populations within specific connectome harmonics results in frequency-specific functional connectivity patterns. This link would imply that, at the whole brain-level, different temporal frequencies support information processing at different spatial scales.

The aim of the present study is to investigate whether connectome harmonics can identify segregation or integration regimes at specific temporal frequencies during sensory perception. To test the validity of this non-invasive approach, we investigated well-known experimental paradigms, in which the neurophysiological processing of information in different frequency-bands has been studied invasively in animal models (4). To this end, we applied connectome harmonics analysis to EEG recordings during an auditory steady-state response (ASSR), which elicits evoked oscillatory activity time-locked to the stimulus onset (23,24), and a visual grating task known to elicit an induced oscillatory response in several frequency bands (25) (Fig. 1a).

Based on the same underlying physical principles underlying the oscillating string, i.e., the wave equation – which states that the frequency of resonance is inversely proportional to the length of the string–, we hypothesized that high-frequency oscillations should be associated with segregation mechanisms (i.e., short-range functional connectivity), whereas lower frequency oscillations should be associated with (long-range) integration. To validate this hypothesis, and its neurophysiological relevance, we further investigated the correlation of estimated neural integration and segregation in the temporal frequency domain participants’ behavioral performance during the visual task.

## Material and Methods

### Recruitment and Assessment of Participants

The participants included in this study were the control subjects form a larger study from the 22q11DS Swiss Cohort (26–28). For this study subjects had to be aged between 17 and 30 years, and should not have had any past or present neurological or psychiatric disease, use of psychotropic medications, psychopathology, learning difficulties, or premature birth. Further exclusion criteria specific to the task were the presence of auditory impairments documented by audiometry screening for the auditory task and a medical history of epilepsy or epileptic seizures for the visual task. Written informed consent was obtained from participants and/or their parents. The study was approved by the cantonal ethics committee for research and conducted according to the Declaration of Helsinki.

### Auditory Steady State Response paradigm

One hundred 40-Hz amplitude-modulated sounds (ripple tones: 1000 Hz carrier tones, duration 2 seconds) and 10 semi-randomly intermixed sounds without amplitude modulation (flat tones at 1000 Hz) were presented binaurally using intra-aural insert earphones (Etymotic Research, Elk Grove Village, Ill.). The interstimulus interval (ISI) was 2 seconds on average (1.5–2.5 seconds, uniform distribution). In order to ensure attentional engagement, participants were asked to detect the flat tones by button press and ignore the 40-Hz ripple tones that entrained the gamma-band responses (23,24).

### Visual paradigm

The visual paradigm consisted of a centrally presented, circular sine wave grating. The circular grating drifted inwards toward the fixation point position and the speed of this contraction increased (velocity step at 2.2 deg/s) at a randomized time point between 750 and 3000 ms after stimulus onset (25,29,30). The experimental protocol comprised of 240 trials divided into three runs of 80 trials. Participants were instructed to press a button as soon as they noticed a speed increase. Stimulus offset was followed by a period of 1000 ms during which subjects were given visual feedback depending on their response. Before beginning the experiment, all the participants underwent a training session with one researcher to be sure that they understood the task. Behavioral measures were calculated as the percentage of correct answers out of the 240 trials and average reaction time.

### EEG data acquisition and pre-processing

EEG data were continuously recorded with a sampling rate of 1000 Hz using a 256-electrodes HydroCel cap (Magstim-EGI) referenced to the vertex (Cz). The impedance was kept below 30 kΩ for all electrodes and below 10 kΩ for the reference and ground electrodes. The preprocessing steps were performed by using the free academic software Cartool (31)(https://sites.google.com/site/cartoolcommunity/home) as previously described (24,30,32,33). In detail, the number of electrodes was reduced from 256 to 204 by eliminating the noisy electrode signals from the cheeks and neck. The data were band-pass filtered between 1 and 140 Hz using non-causal Butterworth filters and we applied additional notch filters at 50, 100, and 150 Hz. The periods of artefacts (e.g., muscle contraction) were marked manually and excluded from further analysis. Noisy channels were identified by means of visual inspection and excluded. Independent Component Analysis was used to remove eye movements (eye blinks and saccades) and ECG artefacts components using a Matlab script based on the EEGlab runica function EEGLAB (https://sccn.ucsd.edu/eeglab/) (34,35). The identified noisy channels were interpolated using a 3D spline interpolation (36). Finally, we applied an instantaneous spatial filter implemented in Cartool that removes local outliers by spatially smoothing the maps without losing their topographical characteristics (37) and data were recalculated to the common average reference.

### MRI data acquisition and pre-processing

We acquired T_1_-weighted images with a 3-T Siemens Prisma at Campus Biotech in Geneva. The parameters for the acquisition of structural images for the T_1_-weighted MPRAGE sequence were as follows: TR=2500 ms, TE=3 ms, flip angle=8°, acquisition matrix=256×256, field of view=23.5 cm, voxel size=0.9×0.9×1.1 mm, and 192 slices. T_1_-weighted images underwent fully automated image processing with Freesurfer, version 6 (38), comprising skull stripping, intensity normalization, reconstruction of the internal and external cortical surface, and parcellation of cortical and subcortical brain regions based on the Desikan-Killiany parcellation (39). Then average measures of volume were extracted from those regions.

### EEG analysis

For the visual task, only epochs with correct behavioral responses were considered for further EEG-analysis (30). Time-frequency (TF) analysis was performed by using Morlet-wavelet transform (frequencies from 2 to 120 HZ, centered on steps of 2 Hz, with adapted resolution according to the FWHM (full width at half maximum) scheme) in MATLAB 2018b as described in previous papers (24,30).

Time epochs from –1.5 to +1.5 seconds relative to the stimulus onset were averaged to event-related spectral perturbation (ERSPs) (40).

For source analysis, the inverse solution (IS) was computed using the academic software Cartool (31) based on individual T1-weighted images preprocessed in Freesurfer. An approximate number of 5000 solution points were distributed in the individually segmented grey matter mask. We used the Locally Spherical Model with Anatomical Constraints (LSMAC) method for the lead field computation that was age-adjusted to reflect differences across age in skull conductivity and thickness (37). A distributed linear inverse solution (Local AutoRegressive Average; LAURA) was employed to compute a transformation matrix from sensor level to inverse solution (41).

The individual Desikan-Killiany parcellations obtained from Freesurfer were natively aligned on the brain of each individual and then used to label the 5000 solution points from the IS model in 84 ROIs covering cortical and subcortical structures. Using the individual IS models, TF decomposition data from the surface were projected to the source space level, reduced as the power, normalized by the baseline period (−1.5 to -0.3 sec) and gathered as the median of each ROI representing the whole brain.

### Consensus structural connectome

In order to perform a fair comparison across subjects, and to obtain a reliable and robust connectivity estimate (11), a consensus structural connectome was computed from a group of 70 healthy young adults (34 females), from an online dataset (42).

The mean age of the participants was 29.7 (range 18.5-59.2 years). All participants were scanned in a 3-Tesla MRI scanner (Trio, Siemens Medical, Germany) with a 32-channel head-coil. The acquisition consisted in a diffusion spectrum imaging (DSI) sequence with 128 diffusion-weighted volumes and a single b0 volume (maximum b-value 8,000 s/mm2), using 2.2×2.2×3.0 mm voxel size. The DSI data reconstruction protocol is described in (43).

A magnetization-prepared rapid acquisition gradient echo (MPRAGE) sequence sensitive to white/gray-matter contrast (with 1 mm in-plane resolution and 1.2 mm slice thickness) was also applied, and Freesurfer (38) and Connectome Mapper 3 (4,5) were used to segment the gray and white matter from the MPRAGE volume and to parcellate the gray-matter with the Desikan-Killiany parcellation (39).

Deterministic streamline tractography on the reconstructed DSI data was performed to estimate the individual structural connectivity matrices, by seeding 32 streamlines per diffusion direction in each white matter voxel. The structural connectome was then defined as the number of fibers connecting each pair of brain regions given by the same parcellation used for the MPRAGE data. Within each brain region, the number of fibers found at the gray matter/white matter interface was summed.

From the connectomes of 70 healthy participants, a group-representative consensus structural brain connectivity matrix was generated as in (11). This method preserves the connection density of each single subject connectome, independently for intra- and interhemispheric connections. By doing so, this method allows more inter-hemispheric connections to be kept in the group estimate, in comparison to simple connectome average across subjects. The connection density of the consensus connectome is approximately 25%.

### Connectome spectral analysis

Connectome spectral analysis explains the brain activity signals in terms of their relationship with structural connectivity. The structural connectivity can be decomposed into Fourier modes, defining the so-called *connectome harmonics* (17). These are the graph equivalent transform to the Fourier decomposition of temporal series (21).

They allow to express brain activity at each time point as a weighted combination of coefficients that define the contribution of each harmonic. Given the parcellated source reconstructed signal in time and frequency (*s*^*t,f*^), we can obtain the contribution of each brain harmonic 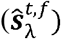 as:

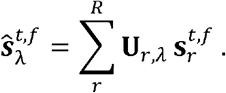

Where **U**_*r,λ*_ indicates the weight of the *λ*-th connectome harmonic at the *r*-th brain region.

### Broadcasting direction

Connectome harmonics are intrinsically ordered according to their spatial frequency in the connectivity graph (from low- to high-frequency harmonics). The broadcasting direction (BD^*t,f*^) is a real-valued index that indicates, for each time-point *t* and temporal frequency *f*, whether the brain activity signal is polarized towards low- or high-frequency connectome harmonics. Specifically, for each time-point we compute:

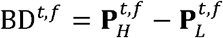

Where 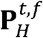 (and 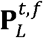) correspond to the proportion of the source reconstructed signal contained within high- (and low-) frequency connectome harmonics:

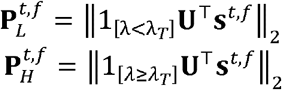

The dichotomization between high and low harmonics (i.e., the selection of *λ*_*T*_) is based on the temporal statistics of the data and is described in the original implementation (20).

Positive (negative) values of BD indicate that the source reconstructed EEG signal is dominated by high-(low-) frequency connectome harmonics, thus more prone to present activity in localized brain regions (large-scale networks).

The significance of the broadcasting direction was assessed by a two-staged statistical test. In the first step, a non-parametric test is performed using permutations and cluster-level correction (46).

A second test is then performed through Monte-Carlo simulations. Specifically, we designed a permutation test to generate a null model distribution of “no effect of structural connectivity on the broadcasting direction”. This null model was generated by computing the broadcasting direction for *n*_*perm*_ connectome surrogates. *n*_*perm*_ was computed taking into account a significance level of α = 0.05, and the number of comparisons to be corrected for by the Bonferroni method, as follows:

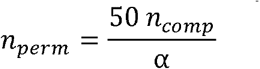

Connectome surrogates consist of randomized graphs obtained using the Brain Connectivity Toolbox (47) function *null_model_und_sign*. The harmonics from these graphs were obtained using the same approach as for the consensus connectome, and they are later referred to as surrogate harmonics in this manuscript.

The estimated broadcasting direction was considered significant if it was within 5% of the extremes of the null model distribution. For this purpose, the uncorrected *p*-value was computed for each time-point as:

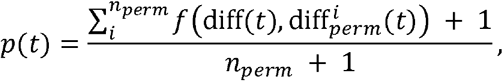

where:

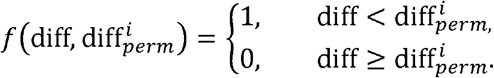

Finally, we corrected the obtained p-values for multiple comparisons with the Benjamini-Hochberg FDR method (48), using the Python toolbox *statsmodels*.

### Joint time-vertex spectral representation

The joint time-vertex spectral representation consists on the simultaneous transformation of the brain activity to both its connectome spectrum and to its temporal spectrum. Here we used the graph Fourier transform for the former and the Morlet-wavelet for the latter transform.

## Results

### Joint time-vertex spectral representation of brain activity

We transformed the source-reconstructed EEG signal, at each time point, jointly by the Morlet-wavelet transform (time domain) and the connectome Fourier transform (brain connectivity domain), yielding a weighed combination of active temporal frequencies and connectome harmonics (see Methods: Joint time-vertex spectral representation).

Our first observation was that this joint time-vertex spectral representation is more compact, as assessed by the *ℓ*_1_/*ℓ*_2_ ratio (median 1075.30, vs. 1525.25 for Morlet Wavelet representation, Wilcoxon signed rank test p-value<.001). This result favors the description of the pattern of brain activation as a specialized process in terms of the involvement of few brain networks, compared to the generalized activation in the brain region representation. We isolated the brain networks underpinning the dynamic responses to the different stimuli observed in Fig. 2b,b’. These brain networks captured (individually) more than of the 30% of the variance of a given oscillatory rhythm.

**Figure 2.**
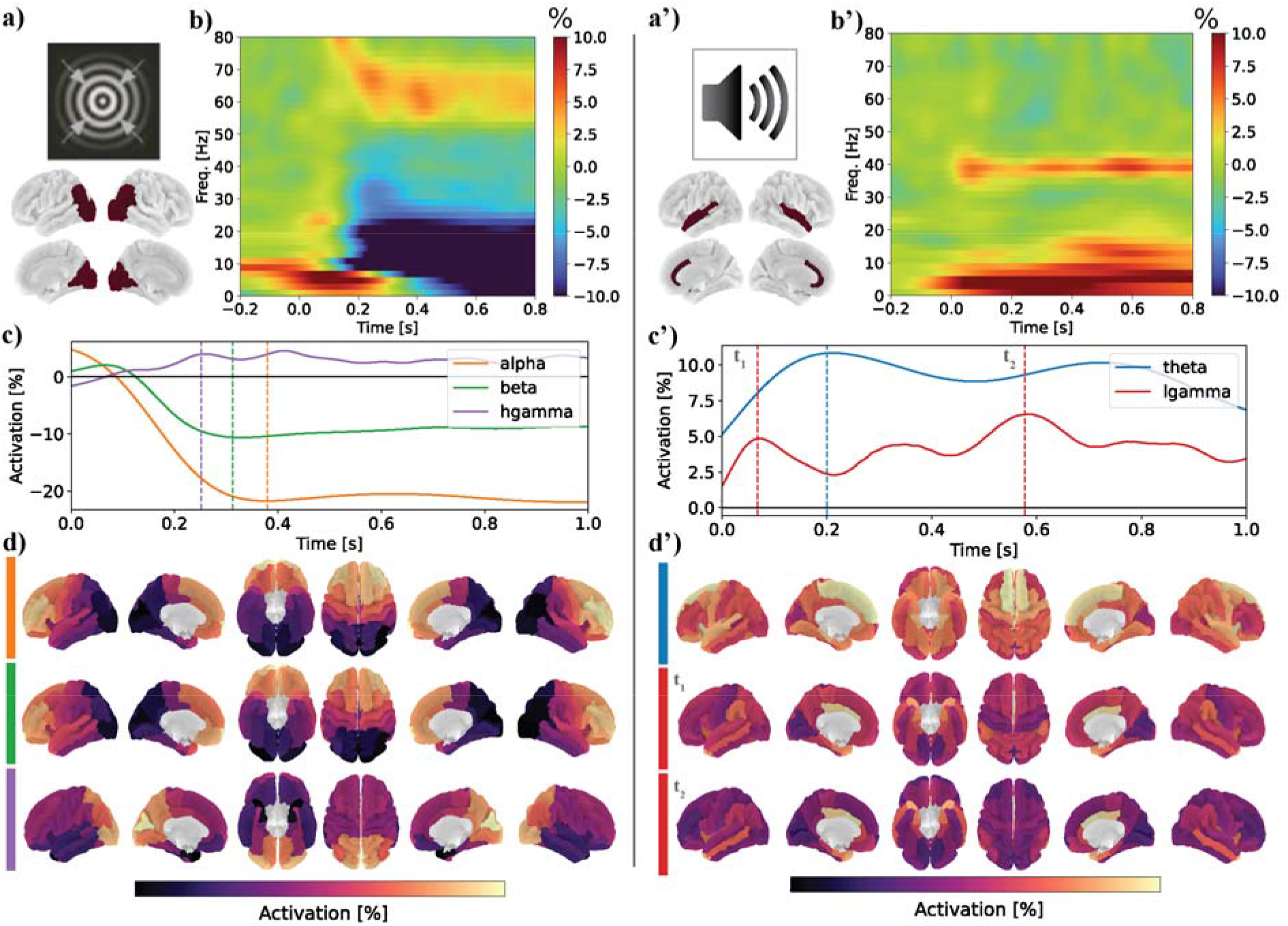
Time-frequency and connectome harmonic representation of brain activity. **a-a’)** Brain regions where the response to the stimulus is most salient: early visual areas for the visual task (a) and temporal regions and the anterior cingulate cortex for the auditory task (a’). **b-b’)** Time-frequency plot of the source reconstructed brain activity during the visual (a) and auditory (a’) task experiments. The average is taken across the brain regions shown in a-a’) and across participants (N=27). **c)** Time series of the average brain activity (for the ROIs shown in a) evoked by the visual stimulus in the relevant frequency bands (alpha = [9, 12], beta = [15, 30] and high gamma = [58, 68] Hz). **c’)** Time series of the average brain activity (for the ROIs shown in a’) in response to the auditory stimulus in the relevant frequency bands (theta = [4, 8] and low gamma = [38, 42] Hz). **d**,**d’)** Reconstruction of the brain activity patterns from the main connectome harmonics (see Methods) at the frequency peaks indicated with dashed lines in c).

In the visual task, at around 0.2 seconds, there is a peak in high gamma activity (58-68 Hz) corresponding to activation of the primary visual cortex and surrounding areas. At about the same time there is trough in the alpha (9-12 Hz) and beta (15-30 Hz) bands reflecting desynchronization (Fig. 2c), corresponding to a wider fronto-parietal network (Fig. 2d).

In the auditory task, oscillatory activation is time-locked to the stimulus onset and corresponds to the activation of temporal regions and the anterior cingulate cortex (ACC) for gamma oscillations (38-42 Hz) and a wider fronto-parieto-temporal network for theta oscillations (4-8 Hz) (Fig. 3d).

**Figure 3.**
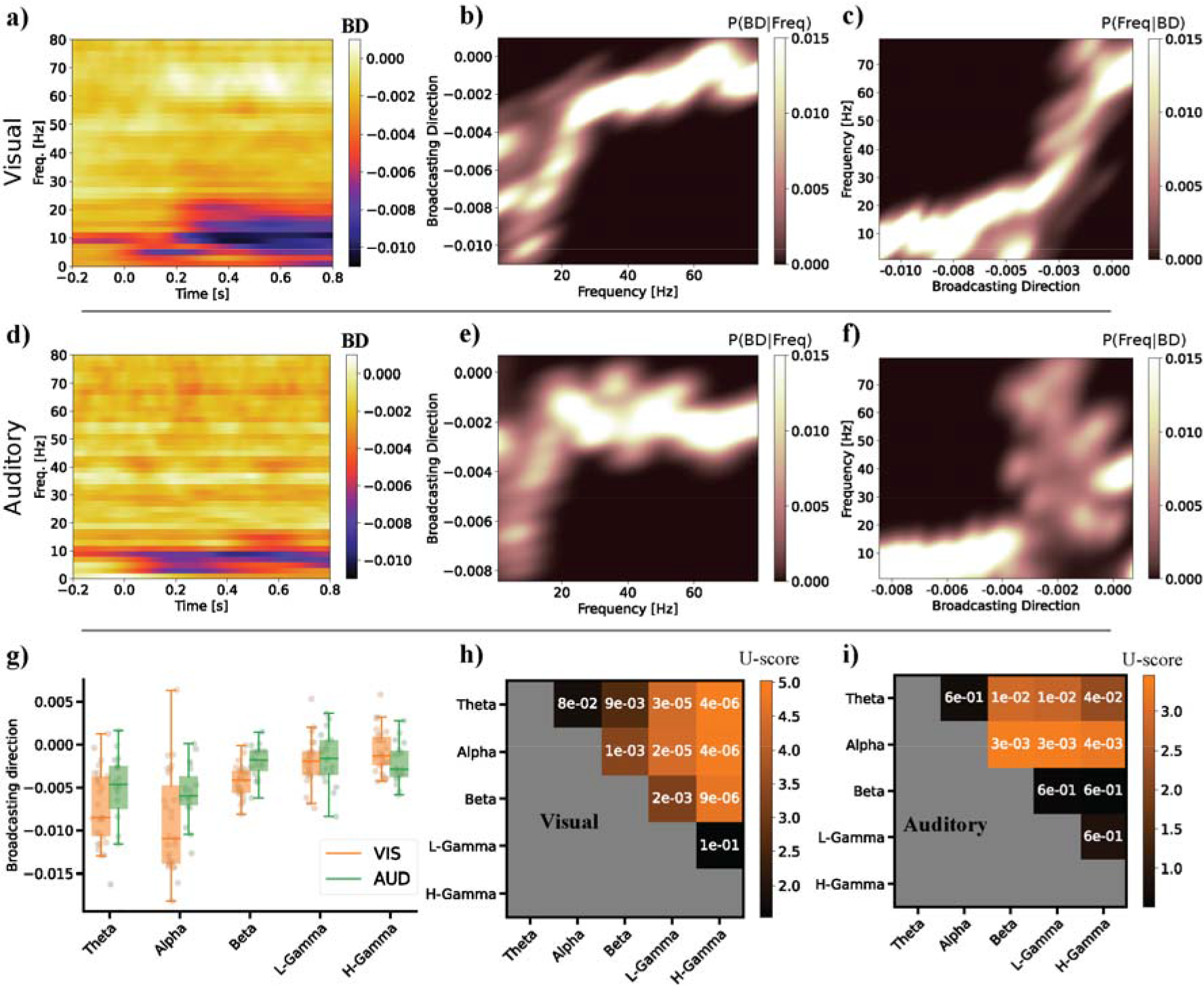
Frequency bands as broadcasting channels. **a)** Frequency-resolved broadcasting direction (BD) for each time-point in both the visual task experiment. More positive BD values are associated to functional segregation mechanisms whereas more negative values indicate tendency towards functional integration. **b)** Probability of a broadcasting direction during the visual task conditioned on the frequency of brain activity. **c)** Probability of the brain activity frequency conditioned on the BD. **d-f)** Same as a-c) for the auditory task. **g)** Average BD for the main frequency channels estimated from the brain activity elicited during both tasks (theta = [4, 8], alpha = [9, 12.5], beta = [15, 30], low-gamma = [38, 42] and high-gamma = [58, 68] Hz). The boxplot distribution shows the variability across participants (N=27 for visual task, N=19 for auditory task). **h)** Statistical testing of the differences of BD among different frequency bands, i.e., distributions shown in g)). The numbers indicate the Ranksum’s test p-value and the color represents the value for the test statistic. **i)** Same as h) for the auditory task.

### Frequency bands as channels for broadcasting brain activity

Next, we computed the broadcasting direction (BD, see *Methods*) of brain activity for each time point and temporal frequency, following the methodology we introduced previously in (20). Briefly, the BD summarizes in a single value the connectome harmonic composition of the source-reconstructed EEG. A positive BD value indicates that the EEG signal pattern has a bias toward connectome harmonics with large eigenvalue (high graph-frequency), which defines an important contribution of the brain networks responsible for the segregation mechanisms. On the contrary, a negative BD value indicates a more important contribution of the integrative functional networks. Fig. 3a,d show how, in both visual and auditory tasks, there is a dominance of integration in the alpha and theta frequency bands (right after stimulus presentation). Furthermore, the segregation mechanisms appear to be specific to higher frequencies, namely high-gamma in the visual task and low gamma in the auditory task.

Fig. 3b,e illustrates how BD varies as a function of frequency band. Specifically, it shows the conditional probability of a BD value given the activation pattern at a frequency in Hz. Brain activation patterns at low frequencies (e.g., up to 30 Hz) tend toward negative BD values, indicating the dominance of integration dynamics due to increased wide-range functional connectivity. At the higher end of the frequency spectrum, activity tends to have a much smaller negative direction, indicating that this frequency range could be used primarily for segregation mechanisms involving more local range functional connectivity.

Based on our results showing the dominance of specific frequency bands in the neural response toward different stimuli (Fig 2), we summarized the BD values for theta, alpha, beta and gamma (low and high) bands. Fig. 3g shows the distribution of BD values across these bands, indicating similar BD specialization in response to visual and auditory tasks. Statistical analysis of the difference between these frequency bands indicates that they mainly cluster into two groups: the first consists of the theta and alpha bands, which are associated with integration mechanisms, whereas the second group consists of beta, low-gamma and high-gamma, and are linked to segregation mechanisms (Wilcoxon ranksum’s test p-value<.05, corrected for multiple comparisons with FDR, see Fig. 3h,i).

### Broadcasting direction predicts behavioral performance

During the visual stimulus task, we recorded the percentage of accurate responses for each participant and reported their distribution in Fig. 4a. We tested whether behavioral responses were related to the BD. For each time point and temporal frequency, we calculated the Spearman’s correlation coefficient between the BD value and the mean behavioral score (detection rates) of all participants (see Fig. 4b). The results show an initial integration in the alpha band during the first 300ms, followed by an increase in segregation (positive BD) in the lower gamma range (Spearman’s correlation test compared against null distribution of shuffled participants’ score, p-value<0.05, corrected for multiple comparisons with Bonferroni method). This contrasts with the more generalized pattern of correlation of the performances with signal power, which partially overlaps (see Fig. 4d).

**Figure 4.**
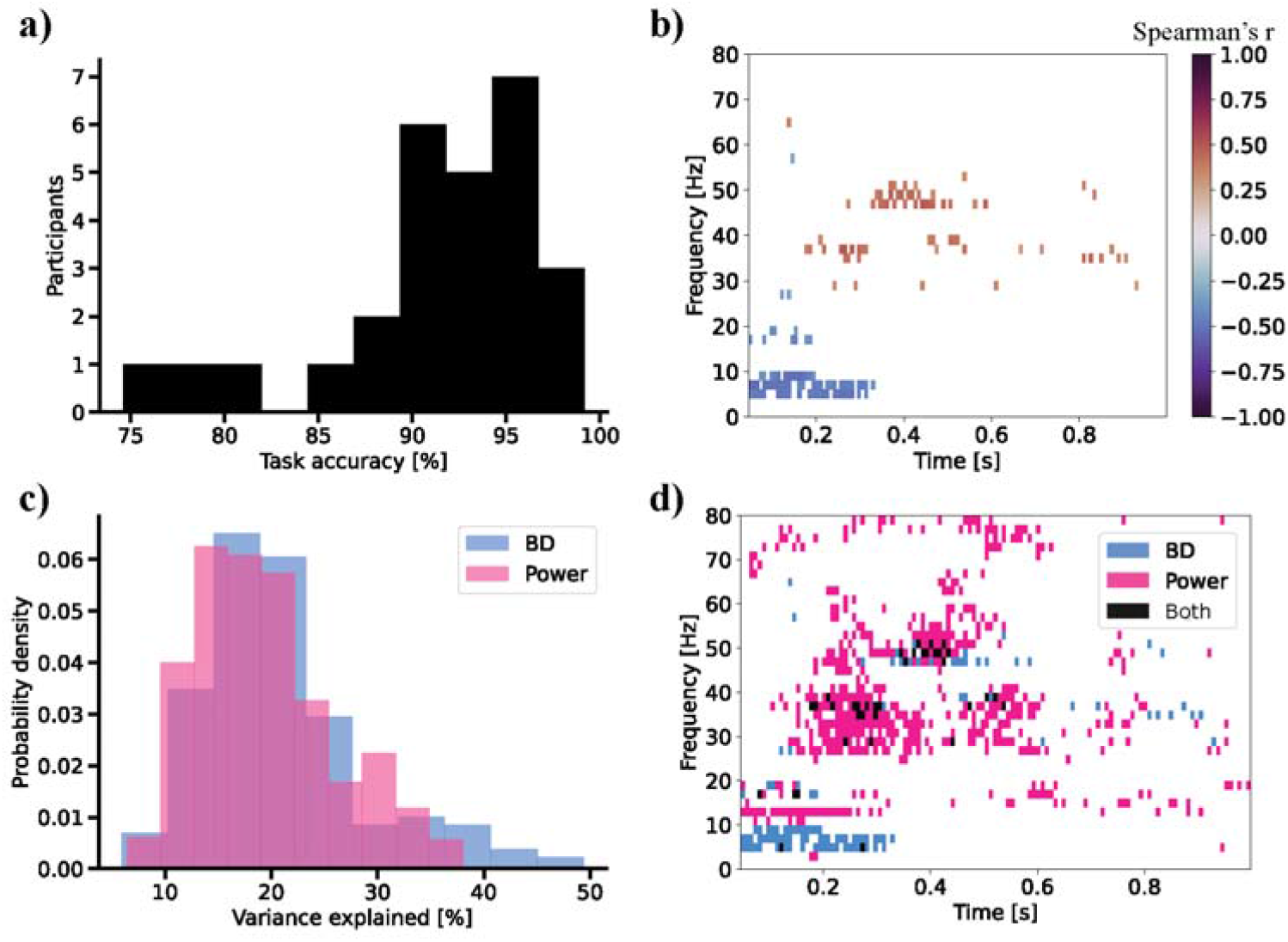
Broadcasting direction predicts behavioral scores. **a)** Distribution of the behavioral scores of the visual task experiment (N=27). The score indicates the task accuracy, i.e. the percentage of correct answers indicating the detection of the visual grating speed change. **b)** Spearman’s correlation between BD and task accuracy, for different frequencies and time-points. Only significant correlation values are shown (after multiple comparisons correction). **c)** Probability densities of the variance of task accuracy explained by the broadcasting direction (blue) and signal power across brain regions. The distribution is defined across all bins showing significant correlation in time and frequency. **d)** Time-frequency plot of the bins in which Spearman’s correlation between task accuracy and BD (blue) and power (pink) are significant. Those time-frequency bins that are significant for both BD and power are colored in black.

## Discussion

Recent efforts to constrain dynamic functional connectivity to anatomical connectivity have shown improved reconstruction of connectivity in electrophysiological signals (49) and fMRI (50). We have introduced and validated with existing literature using intra cortical recordings, a new approach to estimate the frequency-resolved dynamics of different modes of information processing in the brain. Specifically, our method estimates the different modes of information processing by describing the source-reconstructed EEG signal in terms of connectome harmonics (17), i.e., the natural normal modes (spatial frequencies) of the brain connectome (Fig. 1c). Our method naturally incorporates anatomical priors into the estimation of functional connectivity, defining brain activity in a joint time-vertex spectral representation. In this representation, the pattern of brain activity is understood as a weighted combination of the active connectome harmonics at different temporal frequencies.

We have applied this joint time-vertex spectral representation of brain activity on two different datasets, an auditory steady state response (ASSR), which elicits evoked oscillatory activity time-locked to the stimulus onset (23,24), and a visual grating task known to elicit induced oscillatory response in several frequency bands (see Fig. 1). We found that the joint spectral representation has a highly structured pattern (compact), showing specific activation for only a few connectome harmonics at a time, and specific frequencies, compared to the more generalized activation of brain regions (only temporal spectrum). This result indicates that the joint time-vertex spectral representation captures the major axes of variability in macroscopic brain activity data with a smaller number of elements, i.e., their degrees of freedom, as shown by a higher compression of the data.

Importantly, this structured time-vertex spectrum also suggests that brain electrical activity follows similar principles underlying the wave equation (12), i.e., large-scale synchronized networks oscillate at low temporal frequencies, whereas short-range localized networks oscillate fast. Thus, evidence is added to the notion that brain spatiotemporal oscillatory patterns are profoundly shaped by the underlying topology of the connectome. Although this result might be expected in part from the literature showing that functional connectivity networks can be sparsely reconstructed by connectome harmonics (18–20), to our knowledge, this is the first evidence for compactness, in terms of *ℓ*_1_/*ℓ*_2_ ratio, in the join time-vertex spectral domain.

Using the joint-time vertex spectral representation, we isolated the connectome harmonics that underpin the dynamical response to the different tasks (Fig. 3). Looking at the main networks involved in the perceptual processes, we found consistent patterns in the two experiments. The peak gamma activity corresponded to the activation of connectome harmonics that captured the primary sensory cortex and the surrounding areas, i.e., the areas of the early visual system for the visual task, and the temporal regions and the anterior cingulate cortex for the auditory task. In contrast, at lower frequencies, we found connectome harmonics involved in more complex cognitive processes: the fronto-parietal network in the alpha and beta-bands for the visual task, and a fronto-parieto-temporal network in theta for the auditory task. These results also agree with the current literature (51,52) and validate the representation of brain activity in the connectome spectral domain.

After the methodological validation of the joint time-vertex spectral representation, we used it to estimate the broadcasting direction (BD) of the source-reconstructed EEG data. The BD index describes, at a given point in time and frequency range, the connectome harmonic composition of the source-reconstructed EEG signal, indicating the dominance of segregation or integration mechanisms. Inspired by recent developments in the field of graph signal processing (12), our work generalizes the concept of BD dynamics (20) to the temporal frequency domain. A negative BD indicates that low-frequency harmonics are more active than high-frequency harmonics, suggesting a predominance of integration mechanisms over distant brain regions. On the other hand, positive values suggest that segregation mechanisms are mainly at play. Our results show a very strong relationship between temporal frequencies and BD, further supporting the idea that the frequency of synchronization between brain regions is intimately linked to the underlying connectivity structure (Fig. 3).

For both tasks, the BD had its largest values at high and low gamma, indicating that these temporal frequency bands tend toward segregation mechanisms. Theta and alpha show negative values, indicating a tendency toward more integrative mechanisms. While beta band activity has, in general, a balanced composition of connectome harmonics for the visual task, it is associated with slightly more segregation mechanisms in the ASSR task. This close relationship between information processing modes and oscillation frequency is again tightly linked with the wave equation model, as previous studies had suggested (53–56). Using a computational model of brain activity propagated through the structural connectome, the authors of (53) suggest that brain activity could be composed of wave-like patterns that are reconfigured over time. Recent studies further suggest a functional specialization of these brain waves (54,55), as well as a dysfunctional involvement in the development of epileptic seizures (56).

The communication through coherence hypothesis (3,10) suggests that the transmission of information between distant neuronal populations is mediated by different oscillatory frequencies. At the network level, this would translate into networks with long average connectivity (low-frequency harmonics in charge of integration mechanisms) oscillating at low frequency (theta and alpha), and networks with short average connectivity (high-frequency harmonics in charge of segregation mechanisms) oscillating at high-frequency (gamma). This hypothesis has also been supported by results from computational neuroscience models (57), and is in agreement with our results. Overall, our findings suggest that the application of connectome spectral analysis could yield new insights into this theory by identifying the (temporal) spectral composition of each (spatial) harmonic under different experimental conditions or simulations.

Finally, we investigated whether the broadcasting direction (BD) could predict behavior. We found that the performance in the visual task is associated with greater segregation in gamma activity and greater integration in the alpha band range (Fig. 6). These results suggest that the brain state resides in an optimal balance between specialized activity at the level of the visual cortex and integration in prefrontal and associative regions. Alpha desynchronization is thought to mediate top-down information processing, reflecting attentional processes toward visual stimuli (58,59), and thus our findings may reflect the top-down propagation of alpha oscillations across different brain regions to achieve maximal integration. On the other hand, it has been proposed that gamma-band oscillations transmit the propagation of sensory stimuli in the respective sensory cortices (3). Therefore, greater segregation in the gamma band range may indicate greater efficiency in local processing at the level of the visual cortices.

In summary, this study presents a novel tool to explore new hypothesis on brain communication mechanisms linked to the brain connectivity structure. Using this tool, we have provided new insights into the complex interplay between integration and segregation mechanisms in the frequency domain, in the specific cases of visual and auditory perception. Our results suggest that different communication mechanisms take place in different frequency channels broadcasting information at different spatial scales, and that they predict the performance in a behavioral task.

### Limitations and future research

The study of brain activity using connectome harmonics has some limitations related to the fact that diffusion MRI-based tractography contains artefactual connections. However, this is not a crucial limitation since the most important variability in connectome harmonics comes from the local connectivity of the brain (60).

Another limitation is the use of spatial smoothing priors in the source reconstruction algorithm, as this imposes a small but consistent shift of energy towards the low frequency connectomic harmonics (20). Our method of estimating the broadcasting direction aims at being agnostic to this bias by choosing a threshold on the connectome spectrum that divides the energy of the integration and segregation into two equal parts. In this way, if the energy is shifted to one end of the spectrum, the threshold is moved accordingly.

## Acknowledgements

The authors would like to thank the funding agency Swiss National Science Foundation Sinergia for their Grant 170873 to Patric Hagmann, PP00P1_183714, PP00P1_190065 to Gijs Plomp. The authors also would like to thank Mikkel Schöttner and Hugo Fluhr for their useful comments.

## Author contributions

Conceptualization: JRQ, VM, SE, PH; Methodology: JRQ, VM; Software: JRQ, VM; Validation: JRQ, VM; Formal analysis: JRQ, VM, SE, PJU, PH; Investigation: JRQ, VM; Data Curation: VM, VR, CL, CMM; Writing - Original Draft: JRQ, VM; Writing - Review & Editing: All; Visualization: JRQ, VM; Supervision: PH, GP, SE; Project administration: PH, SE; Funding acquisition: PH, GP, CMM, SE

## References

1. Buzsáki G, Draguhn A. Neuronal olscillations in cortical networks. Science (80-). 2004;304(5679):1926–9.

2. Lakatos P, Shah AS, Knuth KH, Ulbert I, Karmos G, Schroeder CE. An oscillatory hierarchy controlling neuronal excitability and stimulus processing in the auditory cortex. Vol. 94, Journal of Neurophysiology. 2005. p. 1904–11.

3. Fries P. Rhythms for Cognition: Communication through Coherence. Neuron [Internet]. 2015;88(1):220–35. Available from: http://dx.doi.org/10.1016/j.neuron.2015.09.034

4. Bastos AM, Vezoli J, Bosman CA, Schoffelen JM, Oostenveld R, Dowdall JR, et al. Visual areas exert feedforward and feedback influences through distinct frequency channels. Neuron. 2015;85(2):390–401.

5. van Kerkoerle T, Self MW, Dagnino B, Gariel-Mathis M-A, Poort J, van der Togt C, et al. Alpha and gamma oscillations characterize feedback and feedforward processing in monkey visual cortex. Proc Natl Acad Sci [Internet]. 2014 Oct 7;111(40):14332–41. Available from: https://pnas.org/doi/full/10.1073/pnas.1402773111

6. Vezoli J, Vinck M, Bosman CA, Bastos AM, Lewis CM, Kennedy H, et al. Brain rhythms define distinct interaction networks with differential dependence on anatomy. Neuron [Internet]. 2021 Dec;109(23):3862-3878.e5. Available from: https://linkinghub.elsevier.com/retrieve/pii/S089662732100725X

7. Kerrén C, Linde-Domingo J, Hanslmayr S, Wimber M. An Optimal Oscillatory Phase for Pattern Reactivation during Memory Retrieval. Curr Biol. 2018;28(21):3383-3392.e6.

8. Shine JM. Neuromodulatory Influences on Integration and Segregation in the Brain. Trends Cogn Sci [Internet]. 2019 Jul;23(7):572–83. Available from: https://linkinghub.elsevier.com/retrieve/pii/S1364661319300944

9. Deco G, Tononi G, Boly M, Kringelbach ML. Rethinking segregation and integration: contributions of whole-brain modelling. Nat Rev Neurosci [Internet]. 2015 Jul 17;16(7):430– Available from: https://www.nature.com/articles/nrn3963

10. Fries P. A mechanism for cognitive dynamics: Neuronal communication through neuronal coherence. Trends Cogn Sci. 2005;9(10):474–80.

11. Betzel RF, Griffa A, Hagmann P, Mišić B. Distance-dependent consensus thresholds for generating group-representative structural brain networks. Netw Neurosci [Internet]. 2019 Jan;3(2):475–96. Available from: https://direct.mit.edu/netn/article/3/2/475-496/2219

12. Grassi F, Loukas A, Perraudin N, Ricaud B. A Time-Vertex Signal Processing Framework: Scalable Processing and Meaningful Representations for Time-Series on Graphs. IEEE Trans Signal Process [Internet]. 2018 Feb 1;66(3):817–29. Available from: http://ieeexplore.ieee.org/document/8115204/

13. Raghavachari S, Lisman JE, Tully M, Madsen JR, Bromfield EB, Kahana MJ. Theta oscillations in human cortex during a working-memory task: Evidence for local generators. Vol. 95, Journal of Neurophysiology. 2006. p. 1630–8.

14. Dickson CT, Biella G, De Curtis M. Evidence for spatial modules mediated by temporal synchronization of carbachol-induced gamma rhythm in medial entorhinal cortex. J Neurosci. 2000;20(20):7846–54.

15. Canolty RT, Soltani M, Dalal SS, Edwards E, Dronkers NF, Nagarajan SS, et al. Spatiotemporal Dynamics of Word Processing in the Human Brain. Front Neurosci. 2007;1(1):185–96.

16. Canolty RT, Knight RT. The functional role of cross-frequency coupling. Trends Cogn Sci. 2010;14(11):506–15.

17. Atasoy S, Donnelly I, Pearson J. Human brain networks function in connectome-specific harmonic waves. Nat Commun [Internet]. 2016 Apr 21;7(1):10340. Available from: http://www.nature.com/articles/ncomms10340

18. Wang R, Lin P, Liu M, Wu Y, Zhou T, Zhou C. Hierarchical Connectome Modes and Critical State Jointly Maximize Human Brain Functional Diversity. Phys Rev Lett [Internet]. 2019 Jul 15;123(3):038301. Available from: https://link.aps.org/doi/10.1103/PhysRevLett.123.038301

19. Glomb K, Rué Queralt J, Pascucci D, Defferrard M, Tourbier S, Carboni M, et al. Connectome spectral analysis to track EEG task dynamics on a subsecond scale. Neuroimage. 2020;221(February).

20. Rué-Queralt J, Glomb K, Pascucci D, Tourbier S, Carboni M, Vulliémoz S, et al. The connectome spectrum as a canonical basis for a sparse representation of fast brain activity. Neuroimage [Internet]. 2021 Dec;244:118611. Available from: https://linkinghub.elsevier.com/retrieve/pii/S1053811921008843

21. Atasoy S, Deco G, Kringelbach ML, Pearson J. Harmonic Brain Modes: A Unifying Framework for Linking Space and Time in Brain Dynamics. Neurosci [Internet]. 2018 Jun 1;24(3):277–93. Available from: http://journals.sagepub.com/doi/10.1177/1073858417728032

22. Tewarie P, Abeysuriya R, Byrne Á, O’Neill GC, Sotiropoulos SN, Brookes MJ, et al. How do spatially distinct frequency specific MEG networks emerge from one underlying structural connectome? The role of the structural eigenmodes. Neuroimage [Internet]. 2019 Feb;186:211–20. Available from: https://linkinghub.elsevier.com/retrieve/pii/S1053811918320603

23. Grent-’t-Jong T, Gajwani R, Gross J, Gumley AI, Krishnadas R, Lawrie SM, et al. 40-Hz Auditory Steady-State Responses Characterize Circuit Dysfunctions and Predict Clinical Outcomes in Clinical-High-Risk Participants: A MEG Study. Biol Psychiatry [Internet]. 2021; Available from: https://doi.org/10.1016/j.biopsych.2021.03.018

24. Mancini V, Rochas V, Seeber M, Roehri N, Rihs T, Ferat V, et al. Aberrant developmental patterns of gamma-band response and long-range communication disruption in youths with 22q11.2 deletion syndrome. Am J Psychiatry. 2022;179(3).

25. Hoogenboom N, Schoffelen JM, Oostenveld R, Parkes LM, Fries P. Localizing human visual gamma-band activity in frequency, time and space. Neuroimage. 2006;29(3):764–73.

26. Schaer M, Debbané M, Bach Cuadra M, Ottet MC, Glaser B, Thiran JP, et al. Deviant trajectories of cortical maturation in 22q11.2 deletion syndrome (22q11DS): A cross-sectional and longitudinal study. Schizophr Res. 2009;115(2–3):182–90.

27. Mancini V, Sandini C, Padula MC, Zöller D, Schneider M, Schaer M, et al. Positive psychotic symptoms are associated with divergent developmental trajectories of hippocampal volume during late adolescence in patients with 22q11DS. Mol Psychiatry [Internet]. 2020;25(11):2844–59. Available from: http://dx.doi.org/10.1038/s41380-019-0443-z

28. Bagautdinova J, Zöller D, Schaer M, Padula MC, Mancini V, Schneider M, et al. Altered cortical thickness development in 22q11.2 deletion syndrome and association with psychotic symptoms. Mol Psychiatry [Internet]. 2021;(October 2020). Available from: http://dx.doi.org/10.1038/s41380-021-01209-8

29. Grent-’t-Jong T, Gajwani R, Gross J, Gumley AI, Krishnadas R, Lawrie SM, et al. Association of magnetoencephalographically measured high-frequency oscillations in visual cortex with circuit dysfunctions in local and large-scale networks during emerging psychosis. JAMA Psychiatry. 2020;77(8):852–62.

30. Mancini V, Rochas V, Seeber M, Grent-’t-Jong T, Rihs TA, Latrèche C, et al. Oscillatory neural signatures of visual perception across developmental stages in individuals with 22q11.2 deletion syndrome. Biol Psychiatry [Internet]. 2022; Available from: https://doi.org/10.1016/j.biopsych.2022.02.961

31. Brunet D, Murray MM, Michel CM. Spatiotemporal analysis of multichannel EEG: CARTOOL. Comput Intell Neurosci. 2011;2011.

32. Cantonas L, Tomescu MI, Biria M, Jan RK, Schneider M, Eliez S, et al. Abnormal development of early auditory processing in 22q11. 2 Deletion Syndrome. Transl Psychiatry. 2019;15–7.

33. Cantonas LM, Mancini V, Rihs TA, Rochas V, Schneider M, Eliez S, et al. Abnormal Auditory Processing and Underlying Structural Changes in 22q11.2 Deletion Syndrome. Schizophr Bull. 2021;47(1):189–96.

34. Makeig S, Jung TP, Bell AJ, Ghahremani D, Sejnowski TJ. Blind separation of auditory event-related brain responses into independent components. Proc Natl Acad Sci U S A. 1997;94(20):10979–84.

35. Delorme A, Makeig S. EEGLAB: An open source toolbox for analysis of single-trial EEG dynamics including independent component analysis. J Neurosci Methods. 2004;134(1):9– 21.

36. Perrin F, Pernier J, Bertrand O. Spherical splines for scalp potential and current density mapping. Electroencephalogr Clin Neurophysiol. 1989;72:184–7.

37. Michel CM, Brunet D. EEG source imaging: A practical review of the analysis steps. Front Neurol. 2019;10(APR).

38. Fischl B. FreeSurfer. Neuroimage. 2012;62(2):774–81.

39. Desikan RS, Se F, Fischl B, Quinn BT, Dickerson BC, Blacker D, et al. An automated labeling system for subdividing the human cerebral cortex on MRI scans into gyral based regions of interest. 2006;31:968–80.

40. Neuper CU, Pfurtscheller G. Event-related dynamics of cortical rhythms_: frequency-specific features and functional correlates. Int J Psychophysiol. 2001;43:41–58.

41. Grave De Peralta Menendez R, Murray MM, Michel CM, Martuzzi R, Gonzalez Andino SL. Electrical neuroimaging based on biophysical constraints. Neuroimage. 2004;21(2):527–39.

42. Griffa A, Aléman-Gomez Y, Hagmann P. Structural and functional connectome from 70 young healthy adults. Zenodo. 2019;

43. Wedeen VJ, Wang RP, Schmahmann JD, Benner T, Tseng WYI, Dai G, et al. Diffusion spectrum magnetic resonance imaging (DSI) tractography of crossing fibers. Neuroimage [Internet]. 2008 Jul;41(4):1267–77. Available from: https://linkinghub.elsevier.com/retrieve/pii/S105381190800253X

44. Daducci A, Gerhard S, Griffa A, Lemkaddem A, Cammoun L, Gigandet X, et al. The Connectome Mapper: An Open-Source Processing Pipeline to Map Connectomes with MRI. Hess CP, editor. PLoS One [Internet]. 2012 Dec 18;7(12):e48121. Available from: https://dx.plos.org/10.1371/journal.pone.0048121

45. Tourbier S, Aleman-Gomez Y, Mullier E, Griffa A, Bach Cuadra M HP. connectomicslab/connectomemapper3: Connectome Mapper v3.0.0-RC4 (Version v3.0.0-RC4). Zenodo. 2020;

46. Maris E, Oostenveld R. Nonparametric statistical testing of EEG- and MEG-data. J Neurosci Methods. 2007;164(1):177–90.

47. Rubinov M, Sporns O. Complex network measures of brain connectivity: Uses and interpretations. Neuroimage [Internet]. 2010 Sep;52(3):1059–69. Available from: https://linkinghub.elsevier.com/retrieve/pii/S105381190901074X

48. Benjamini Y, Hochberg Y. Controlling the False Discovery Rate: A Practical and Powerful Approach to Multiple Testing. R Stat Soc. 1995;57(1):289–300.

49. Pascucci D, Rubega M, Rué-Queralt J, Tourbier S, Hagmann P, Plomp G. Structure supports function: Informing directed and dynamic functional connectivity with anatomical priors. Netw Neurosci [Internet]. 2022 Jan 26;1–19. Available from: https://direct.mit.edu/netn/article/doi/10.1162/netn_a_00218/108678/Structure-supports-function-Informing-directed-and

50. Crimi A, Dodero L, Sambataro F, Murino V, Sona D. Structurally constrained effective brain connectivity. Neuroimage [Internet]. 2021 Oct;239:118288. Available from: https://linkinghub.elsevier.com/retrieve/pii/S1053811921005644

51. Pagnotta MF, Pascucci D, Plomp G. Selective attention involves a feature-specific sequential release from inhibitory gating. Neuroimage [Internet]. 2022 Feb;246:118782. Available from: https://linkinghub.elsevier.com/retrieve/pii/S1053811921010545

52. Costa-Faidella J, Sussman ES, Escera C. Selective entrainment of brain oscillations drives auditory perceptual organization. Neuroimage [Internet]. 2017 Oct;159:195–206. Available from: https://linkinghub.elsevier.com/retrieve/pii/S1053811917306249

53. Roberts JA, Gollo LL, Abeysuriya RG, Roberts G, Mitchell PB, Woolrich MW, et al. Metastable brain waves. Nat Commun [Internet]. 2019 Dec 5;10(1):1056. Available from: http://www.nature.com/articles/s41467-019-08999-0

54. Raut R V., Snyder AZ, Raichle ME. Hierarchical dynamics as a macroscopic organizing principle of the human brain. Proc Natl Acad Sci [Internet]. 2020 Aug 25;117(34):20890–7. Available from: https://pnas.org/doi/full/10.1073/pnas.2003383117

55. Huntenburg JM, Bazin P-L, Margulies DS. Large-Scale Gradients in Human Cortical Organization. Trends Cogn Sci [Internet]. 2018 Jan;22(1):21–31. Available from: https://linkinghub.elsevier.com/retrieve/pii/S1364661317302401

56. Breakspear M, Roberts JA, Terry JR, Rodrigues S, Mahant N, Robinson PA. A Unifying Explanation of Primary Generalized Seizures Through Nonlinear Brain Modeling and Bifurcation Analysis. Cereb Cortex [Internet]. 2006 Sep 1;16(9):1296–313. Available from: http://academic.oup.com/cercor/article/16/9/1296/276340/A-Unifying-Explanation-of-Primary-Generalized

57. Raj A, Cai C, Xie X, Palacios E, Owen J, Mukherjee P, et al. Spectral graph theory of brain oscillations. Hum Brain Mapp [Internet]. 2020 Aug 23;41(11):2980–98. Available from: https://onlinelibrary.wiley.com/doi/10.1002/hbm.24991

58. Rihs TA, Michel CM, Thut G. A bias for posterior α-band power suppression versus enhancement during shifting versus maintenance of spatial attention. Neuroimage [Internet]. 2009;44(1):190–9. Available from: http://dx.doi.org/10.1016/j.neuroimage.2008.08.022

59. Romei V, Rihs T, Brodbeck V, Thut G. Resting electroencephalogram alpha-power over posterior sites indexes baseline visual cortex excitability. Neuroreport. 2008;19(2):203–8.

60. Naze S, Proix T, Atasoy S, Kozloski JR. Robustness of connectome harmonics to local gray matter and long-range white matter connectivity changes. Neuroimage [Internet]. 2021 Jan;224:117364. Available from: https://linkinghub.elsevier.com/retrieve/pii/S1053811920308508

